# Bifurcated monocyte states are predictive of mortality in severe COVID-19

**DOI:** 10.1101/2021.02.10.430499

**Authors:** Anthony R. Cillo, Ashwin Somasundaram, Feng Shan, Carly Cardello, Creg J. Workman, Georgios D. Kitsios, Ayana Ruffin, Sheryl Kunning, Caleb Lampenfeld, Sayali Onkar, Stephanie Grebinoski, Gaurav Deshmukh, Barbara Methe, Chang Liu, Sham Nambulli, Lawrence Andrews, W. Paul Duprex, Alok V. Joglekar, Panayiotis V. Benos, Prabir Ray, Anuradha Ray, Bryan J. McVerry, Yingze Zhang, Janet S. Lee, Jishnu Das, Harinder Singh, Alison Morris, Tullia C. Bruno, Dario A.A. Vignali

## Abstract

Coronavirus disease 2019 (COVID-19) caused by SARS-CoV-2 infection presents with varied clinical manifestations^1^, ranging from mild symptoms to acute respiratory distress syndrome (ARDS) with high mortality^2,3^. Despite extensive analyses, there remains an urgent need to delineate immune cell states that contribute to mortality in severe COVID-19. We performed high-dimensional cellular and molecular profiling of blood and respiratory samples from critically ill COVID-19 patients to define immune cell genomic states that are predictive of outcome in severe COVID-19 disease. Critically ill patients admitted to the intensive care unit (ICU) manifested increased frequencies of inflammatory monocytes and plasmablasts that were also associated with ARDS not due to COVID-19. Single-cell RNAseq (scRNAseq)-based deconvolution of genomic states of peripheral immune cells revealed distinct gene modules that were associated with COVID-19 outcome. Notably, monocytes exhibited bifurcated genomic states, with expression of a cytokine gene module exemplified by *CCL4* (MIP-1β) associated with survival and an interferon signaling module associated with death. These gene modules were correlated with higher levels of MIP-1β and CXCL10 levels in plasma, respectively. Monocytes expressing genes reflective of these divergent modules were also detectable in endotracheal aspirates. Machine learning algorithms identified the distinctive monocyte modules as part of a multivariate peripheral immune system state that was predictive of COVID-19 mortality. Follow-up analysis of the monocyte modules on ICU day 5 was consistent with bifurcated states that correlated with distinct inflammatory cytokines. Our data suggests a pivotal role for monocytes and their specific inflammatory genomic states in contributing to mortality in life-threatening COVID-19 disease and may facilitate discovery of new diagnostics and therapeutics.

## Introduction

The emergence of SARS-CoV-2 has led to over 104 million cases of COVID-19 worldwide, with more than 2.2 million global deaths^4^. In the United States alone, more than 26 million cases and over 450,000 deaths have been reported^4^. Despite the rapid development and deployment of vaccines^5,6^, thousands of patients are currently hospitalized with severe COVID-19. COVID-19 is also projected to be a major cause of critical illness and hospitalizations throughout 2021 and beyond, before world-wide vaccine campaigns can control the pandemic. Furthermore, the recent emergence of more highly transmissible viral variants (i.e. B.1.1.7 in the United Kingdom and N501Y in South Africa) is predicted to increase infections and hospitalizations in the coming months^7^. Among initial clinical trials examining several pharmacologic interventions in severe COVID-19, non-specific immunosuppression with corticosteroids has shown clear efficacy^8–10^, and recent work has shown efficacy of IL-6 blockade in patients treated soon after ICU admission^11^. Therefore, severe COVID-19 continues to carry a high mortality risk. Thus, in depth analyses of cellular and molecular states, particularly of the immune system, that are associated with survival or death in severe COVID-19 disease are urgently needed. Such analyses will accelerate the development of new diagnostics and therapeutics that could save lives.

SARS-CoV-2 infection can cause acute lung injury leading to ARDS^1^, dysregulated inflammation^12^ and hyper-coagulability^13^. Studies dissecting the immunopathology of COVID-19 have evaluated patients across a spectrum of disease severity (mild to critical), and noted lymphopenia^1^ accompanied with high levels of multiple inflammatory cytokines, CRP^14^ and D-dimer^13^ in the context of severe disease. In particular, elevated levels of the cytokines IL-6, TNFα, IP-10 (CXCL10), IL-8 (CXCL8) and IL-10 have been reported in severe COVID-19 disease^15,16^. Notably, the dysregulated levels of IL-6 in COVID-19 patients are lower compared with those in non-COVID ARDS, cytokine release syndrome or sepsis, suggesting differences in the underlying etiology^17^. Type I interferon (IFN) signaling is essential in moderating COVID-19 disease, as patients with either IFN auto-antibodies^18^ or inborn errors of type I interferon^19^ production have a much higher risk of severe COVID-19. Paradoxically, a delayed and excessive type I IFN response is implicated in severe disease and leads to mortality in a mouse model of SARS-CoV-2 infection^20–22^. To date, no molecular pathways or components that can accurately predict mortality in severe COVID-19 patients (i.e. patients in the ICU) have been described. Thus, we undertook high-dimensional cellular and molecular analyses of the immune system in critically ill ICU patients, utilizing flow cytometry, scRNAseq and cytokine profiling and coupled them with machine learning to reveal divergent cellular states and gene modules that predicted mortality.

## Results

### Clinical cohorts, subject characteristics and study design

Following informed consent, we enrolled 41 consecutive critically ill patients with acute hypoxemic respiratory failure and symptoms suggestive of COVID-19 in a prospective, observational cohort study, with limited longitudinal sampling. Based on reference-standard nasopharyngeal swab SARS-CoV-2 qPCR, COVID-19 was diagnosed in 35 patients (COVID-19 group), whereas a non-COVID etiology of acute respiratory illness was identified in 6 patients who had negative SARS-CoV-2 qPCR (non-COVID ARDS group). Clinical details of this cohort have recently been described in detail^23^. As controls in the study, we also included 10 healthy blood donors (healthy donor group). The median ages of non-COVID ARDS and COVID-19 patients were 62 and 65 years, and many patients had pre-existing conditions (Table S1) including diabetes (17% non-COVID ARDS, 47% COVID-19) and either current or former smoking (67% non-COVID ARDS, 60% COVID-19). Mortality at 90 days was 33% and 40% in non-COVID ARDS and COVID-19 patients, respectively (Table S1). Comparisons of clinical covariates by death or survival (day 90) showed trends towards higher mortality in older patients and patients with lower BMI. We note that patients who received glucocorticoids were more likely to survive (Table S2). We obtained peripheral blood as well as endotracheal aspirates from COVID-19 patients to perform high-dimensional profiling of cellular genomic states and cytokines. Only blood samples were obtained from non-COVID ARDS and healthy donors (Fig. 1a, Methods). We leveraged flow cytometry, scRNAseq and cytokine datasets from blood samples to thoroughly interrogate immune system states across these clinical groups (Fig. 1a, Table S3).

**Figure 1.**
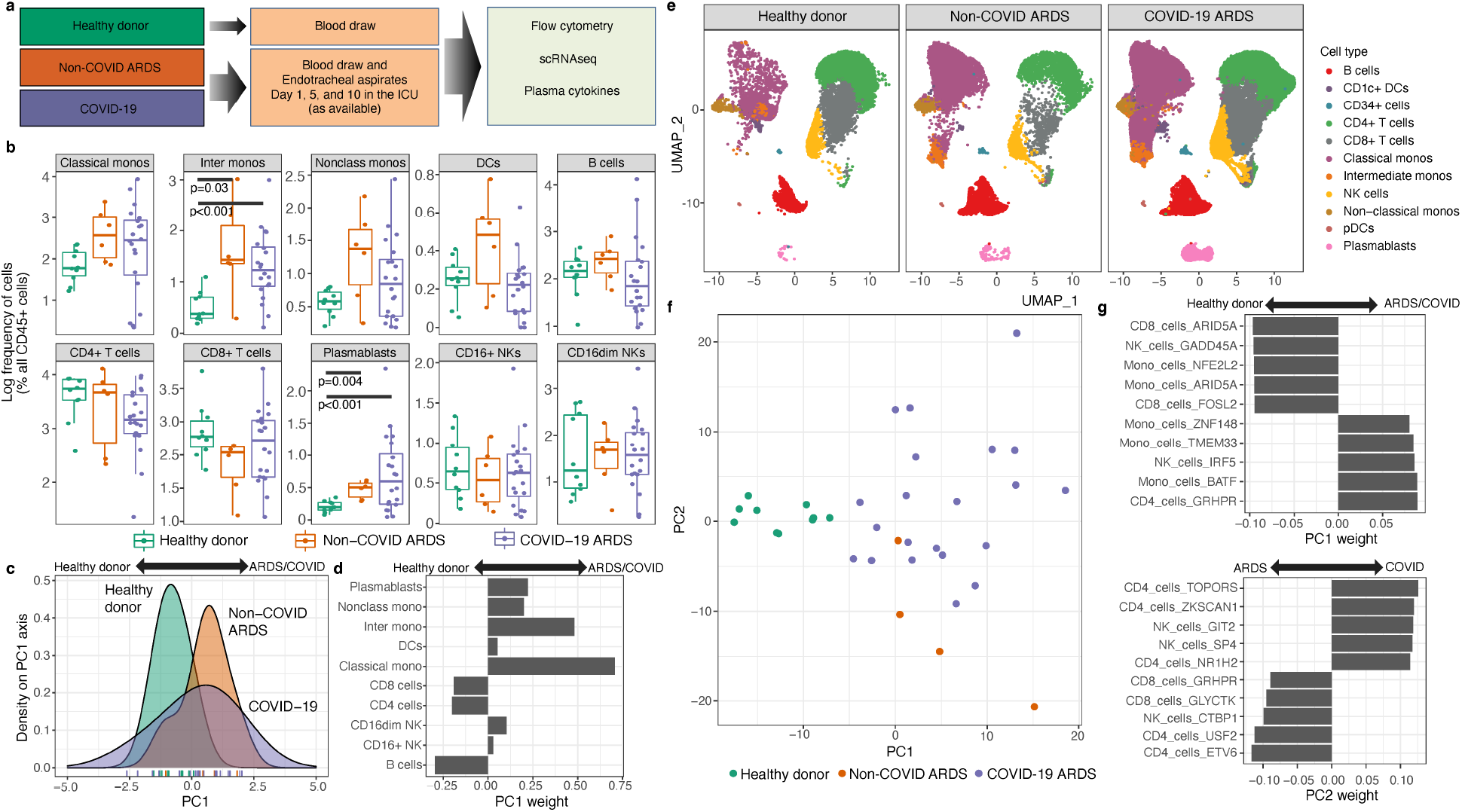
Expansion of intermediate monocytes and plasmablasts is associated with COVID-19. a) Schema highlighting the 3 clinical groups along with biological specimens obtained and assays performed. b) Flow cytometric analysis of immune cell frequencies in peripheral blood of healthy donors (one timepoint), non-COVID ARDS patients (day 1 ICU), and COVID-19 patients (day 1 ICU). c) Principal component analysis (PCA) of immune cell frequencies displayed in panel (b). d) Weightings of immune cell frequencies that contribute to the PC1 embeddings in panel (c) and distinguish critically ill patients (non-COVID ARDS/COVID-19) versus healthy donors. e) Single-cell RNAseq analysis of 98,327 cells showing the canonical immune lineages from peripheral blood of healthy donors (14,271 cells), non-COVID ARDS patients (13,060 cells) and COVID-19 patients (78,922 cells). f) PCA performed by using differentially expressed immune cell gene modules (delineated with Arboreto, see Methods), as molecular features of study participants in the three clinical groups. g) Weightings of the immune cell gene modules that are dominant contributors to PC1 and PC2 embeddings in panel (f) and distinguish critically ill non-COVID ARDS or COVID-19 patients from healthy donors.

### Expansion of intermediate monocytes and plasmablasts in COVID-19

We utilized high-dimensional flow cytometry to evaluate peripheral blood mononuclear cells (PBMC) of critically ill patients (COVID-19 and non-COVID ARDS) at post-enrollment day 1 in the ICU (Fig. 1b and Fig. S1a-c). Overall, we found that at day 1, COVID-19 and non-COVID ARDS patients had significantly higher levels of intermediate monocytes and plasmablasts versus healthy controls. We did not find significant associations between cell frequencies and death or survival (Fig. S2a-b). Although cycling CD8+ T cells that co-express HLA-DR and CD38 are associated with viral infection in general and COVID-19 in particular^24^, there were no differences in frequencies of this CD8+ T cell subset on day 1 between deceased and surviving patients in our cohort (Fig. S2c-d). We performed principal component analysis (PCA) using the immune cell frequencies (from Fig. 1b) to determine whether these multi-variate features could stratify patients by clinical group. When visualized along the PC1 axis, distributions of patient samples demonstrated considerable heterogeneity (Fig. 1c). However, COVID-19 as well as non-COVID ARDS samples were shifted rightwards on PC1, while healthy donors were shifted leftwards. Consistent with analysis in Fig. 1b, higher frequencies of myeloid cells and plasmablasts contributed to the PC1 distribution of samples from critically ill patients, while higher frequencies of T and B cells distinguished the healthy donors (Fig. 1d). Although assessment of cell frequencies revealed shifts in the composition of immune cells in COVID-19 and non-COVID ARDS patients versus healthy donors, these frequencies yielded no statistically significant associations between COVID-19 and non-COVID ARDS or in COVID-19 patient outcome.

### Dynamic gene modules derived from scRNAseq stratify subjects by clinical group

To query the dynamic genomic states of immune cells and generate finer-grained predictive features, we performed scRNAseq analysis of PBMC from COVID-19 and non-COVID ARDS patients and healthy donors (Methods). Samples were obtained from critically ill patients on days 1, 5 and 10 post-enrollment whereas those from healthy donors were obtained at one timepoint (Table S3). In total, we profiled 99,618 cells (Fig. 1e). Canonical immune cell types were identified as previously described(Methods)^25^, and were visualized across clinical groups. In agreement with the flow cytometric analysis, there were increased frequencies of intermediate monocytes and plasmablasts. Qualitatively, there appeared to be distinguishable transcriptional states of classical monocytes in COVID-19 and non-COVID ARDS patients versus healthy donors reflected by the unique area occupied by classical monocytes in the uniform manifold approximation and projections (UMAPs)^26^.

We next sought to determine whether the aforementioned clinical groups could be stratified by the transcriptional states of their immune cells. To analyze transcriptional states across immune cell types, we used Arboreto within the pySCENIC framework^27^(Methods). First, we bioinformatically isolated all major canonical immune cell types across all samples and timepoints (Fig. S3a-b) and then utilized tree-based regression analysis in Arboreto to identify modules of co-expressed genes presumptively co-regulated by given transcription factors (Fig. S3c). Each transcription factor-associated gene module was linked with a cell type, and individual genes within a module were assigned a weight derived from the tree-based regression analysis (Supp. Data S1). Finally, we scored each gene module across all of its linked cells and assigned each patient sample a single score for that module based on its median module score across all linked cells of that patient at a given timepoint (Fig. S3c). The immune cell type linked gene module scores are a reflection of dynamic transcriptomic states of such cells. We then asked whether we could use the resultant gene module activity matrix generated from patient scRNAseq datasets to distinguish between the study groups. To do this, we first identified the top 10 most significantly different gene modules for each immune cell type between the patient groups (Supp. Data S2). We then performed PCA with these selected gene modules and visualized the top 2 PCs. This analysis revealed that PC1 stratified patients by critical illness and that PC2 partially stratified COVID-19 patients from non-COVID ARDS (Fig. 1f). Interestingly, 3 of the top 5 gene modules that enabled separation of COVID-19 and non-COVID ARDS patients from healthy controls were linked to monocytes (Fig. 1g), raising the possibility that monocytic activation states may play an important role in critical illness and outcomes.

### Divergent monocyte activation states in severe COVID-19

After using day 1 gene module scores to stratify patients by clinical group, we next queried whether these cellular and molecular features were associated with COVID-19 patient outcome. To address this question, we first performed a Wilcoxon rank sum test to determine whether gene module scores were significantly associated with mortality. We found a total of 56 statistically significant gene modules (Supp. Data S3), and when we visualized these gene modules across all patients and performed co-clustering analysis, we observed a demarcation between patients who survived and those who did not (Fig. 2a). Notably, we identified several monocyte gene modules that were associated either with survival (e.g., Mono_cells_PDE6H) or with death (e.g., Mono_cells_NOC2L).

**Figure 2.**
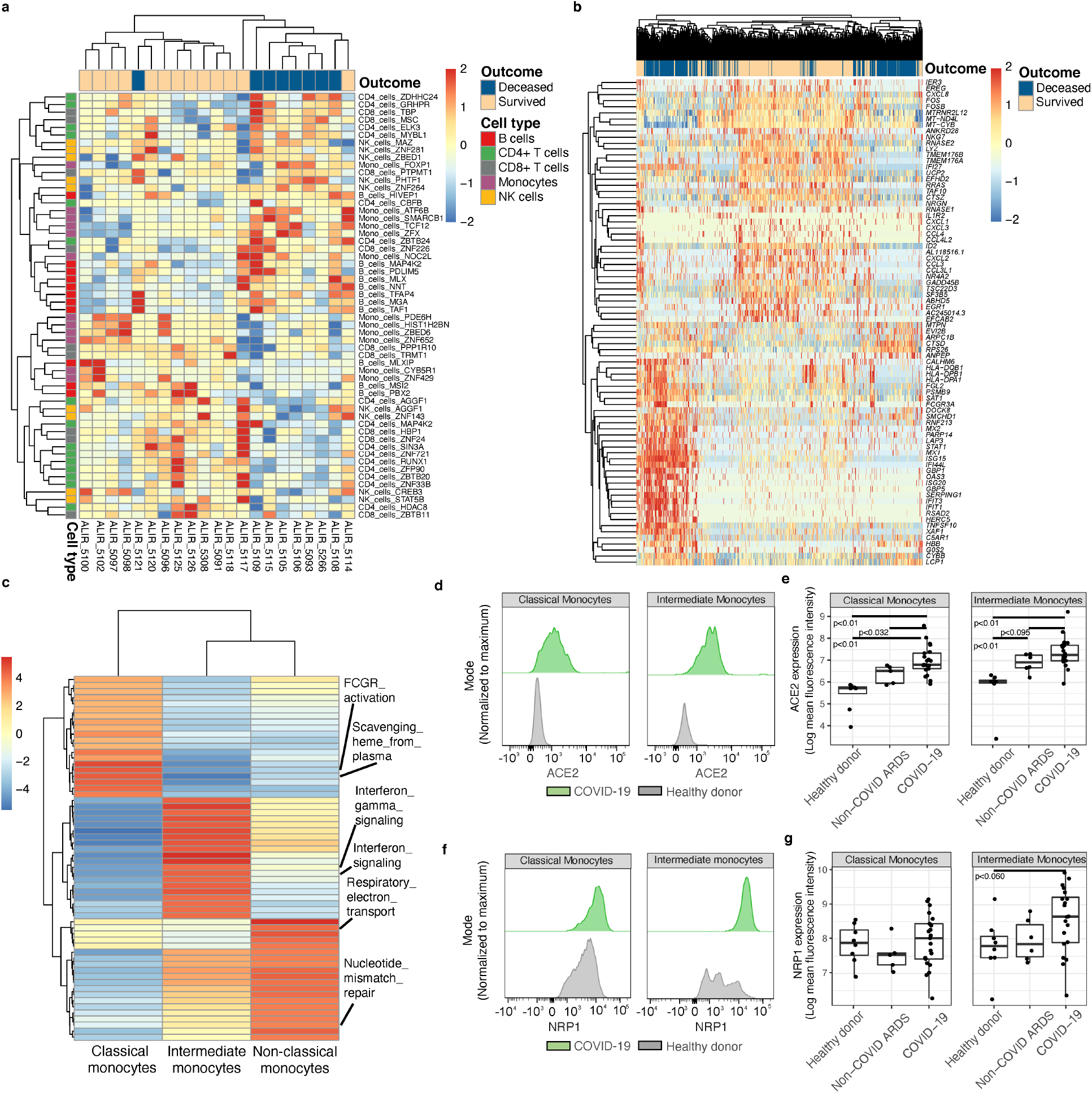
Bifurcated monocyte activation states in severe COVID-19. a) Heatmap displaying hierarchical clustering of Arboreto delineated gene modules that were statistically significantly associated with COVID-19 outcome. Patient samples (day 1 ICU) are arrayed in columns and gene modules in rows. Color bar on the left indicates linkage of gene modules with particular immune cell types. b) Heatmap displaying hierarchical clustering of differentially expressed genes associated with outcome from scRNAseq of peripheral blood monocytes from COVID-19 patients (day 1 ICU). Cells from patient samples with the indicated outcomes are arrayed in columns and genes are annotated in rows. c) Gene set enrichment analysis of canonical monocyte subsets in the scRNAseq dataset. d) Representative flow plots of ACE2 expression on classical and intermediate monocytes. e) Expression levels (mean fluorescence intensity, MFI), on classical and intermediate monocytes in indicated patient groups versus healthy donors. f) Representative flow plots of NRP1 expression on classical and intermediate monocytes. g) Expression levels of NRP1 (MFI) on classical and intermediate monocytes in indicated patient groups versus healthy donors.

To further explore these divergent monocytic states in COVID-19 patients, we performed an orthogonal analysis based on differentially expressed genes (DEGs) associated with death versus survival across all monocyte subsets (Fig. 2b). Analysis of DEGs revealed that 9 of the top 20 genes associated with survival were derived from the Mono_cells_PDE6H module. These genes included the chemokines *CXCL1*, *CXCL3*, *CCL4* (MIP-1β), *CXCL2* and *CCL3* (MIP-1α). Conversely, a set of interferon response genes (*MX2*, *MX1*, *ISG15*, *IFI44L*, *OAS3*, *ISG20*, *IFIT3*, and *IFIT1*) were associated with death in COVID-19 patients. Although these genes were not part of the Mono_cells_NOC2L module, the latter contained genes that encoded components of viral sensing and the RIG-I pathway (Supp. Data S1). Thus, the analyses of monocytic gene modules and DEGs complemented one another while expanding the gene features that distinguished the divergent monocytic states.

To distinguish the activation states of canonical monocyte subsets, we performed gene set enrichment analysis^25^ (Fig. 2c, Supp. Data S4). Gene sets for interferon signaling were highly enriched in intermediate monocytes and suggestive of viral entry into such cells. To explore this possibility, we utilized flow cytometry to analyze the expression levels of the SARS-CoV-2 entry receptor ACE2 and cofactor neuropilin-1 (NRP1). ACE2 expression was significantly higher on both classical and intermediate monocytes in COVID-19 as well as non-COVID-19 ARDS patients compared to healthy donors (Fig. 2d-e). Notably only intermediate monocytes from COVID-19 patients expressed NRP1 at significantly higher levels versus healthy controls (Fig. 2f-g). Expression of these two entry receptors suggests activated monocytes are permissive for SARS-CoV-2 entry. However, we were unable to detect SARS-CoV-2 transcripts in these monocytic cells by scRNAseq (Methods). Consistent with a systemic inflammatory state, both intermediate and classical monocytes expressed higher levels of the chemokine receptor CCR5 (Fig. S4a-b). Thus, our analysis of gene module scores on baseline samples stratified COVID-19 patients based on outcome. In particular, genes involved in interferon signaling in monocytes were associated with mortality.

### Distinctive monocyte inflammatory states in COVID-19 patient lungs

Upon uncovering the distinctive inflammatory gene modules in monocytes that were associated with outcomes, we sought to determine if they were reflective of discrete cell states (canonical or non-canonical) within the heterogeneous monocytic compartment. To achieve this, we re-clustered all PBMC monocytes from all samples, and visualized these cells using either canonical monocyte markers (Fig. 3a), or by unsupervised clustering (Fig. 3b). Cluster 4 represented intermediate monocytes, while clusters 1, 2, 3, 7 and 8 were classical monocytes with cluster 6 representing non-classical monocytes. Next, we analyzed the association between the monocyte clusters and differentially expressed genes associated with outcome. Based on the aforementioned gene modules, we focused on key cytokine and interferon response genes. This analysis mapped the expression of *CCL3*, *CCL4*, *CXCL1*, *CXCL2* and *CXCL3* to the classical monocyte clusters 7 and 8, while the interferon response genes such as *IFI44L*, *MX1*, *MX2, OAS3* and others mapped to intermediate monocyte cluster 4 (Fig. 3c). These findings suggest that a distinctive chemokine programmed state of classical monocytes in the ICU is associated with eventual survival, while high levels of interferon response in intermediate monocytes is associated with death.

**Figure 3.**
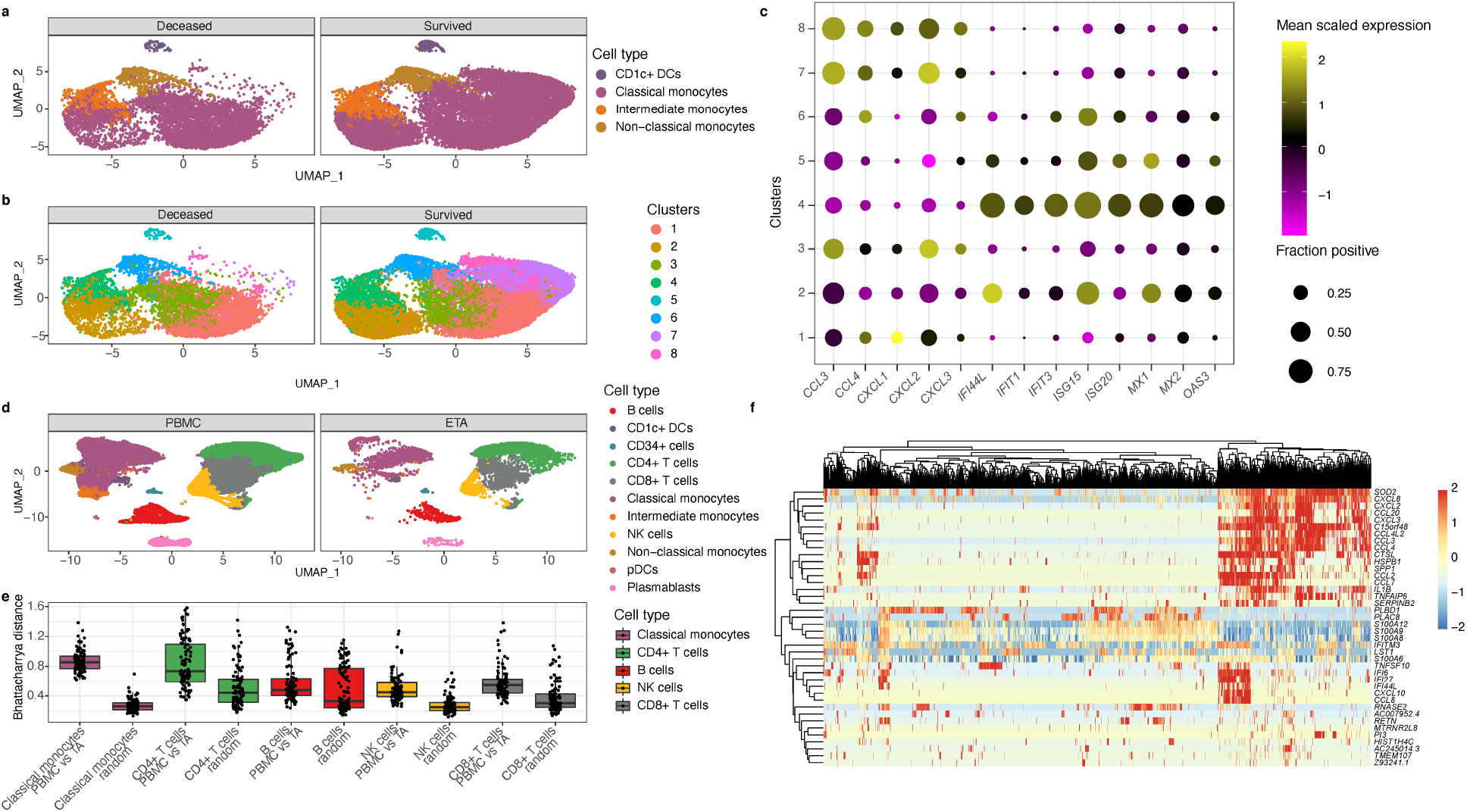
Bifurcated monocytes inflammatory states map to distinct subsets and are reflected in lungs of COVID-19 patients. a) UMAPs showing canonical monocytes clusters in peripheral blood of patients who died versus those who survived. b) Same UMAPs as panel (a) but monocytes are delineated using graph-based clustering. c) Frequency and magnitude of expression of genes reflective of bifurcated inflammatory monocytes states. The frequency of cells expressing the indicated gene (size of the dot) and its mean expression level in that cluster (color of the dot). d) UMAPs of immune cells in COVID-19 patient samples derived from blood or endotracheal aspirates (ETA). e) Plot of Bhattacharyya distances, a similarity measure of distributions of cellular gene expression states, of the indicated immune cells in blood versus ETA of COVID-19 patients. f) Heatmap displaying hierarchical clustering of single-cell expression profiles of monocytes from COVID-19 patient ETA samples. Cells from patient samples are arrayed in columns and annotated genes that were differentially expressed between PBMC and ETA monocytes are in rows.

To determine if these distinctive inflammatory monocytic states revealed in PBMC were also reflective of cells within patients’ lungs in severe COVID-19 disease, we analyzed endotracheal aspirate (ETA) samples from mechanically ventilated patients by scRNAseq. UMAP visualization of ETA samples in comparison with PBMC showed the major immune cell clusters to be largely overlapping, with some distinct regions reflecting enrichment of cells within ETA samples (Fig. 3d). To quantify the degree of difference in gene expression space between corresponding immune cell clusters in matched ETA and PBMC samples, we utilized a high dimensional distance metric known as Bhattacharyya distance (BD; Fig. 3e). A larger BD corresponds to a more dissimilar transcriptional state. Based on this approach, classical monocytes were the most different between ETA and PBMC (Fig. 3e). We were unable to compare the states of intermediate or non-classical monocytes in the ETA samples with this approach, due to limited numbers of these cells. We next performed DEG analysis of classical monocytes from PBMC versus ETA samples to determine if genes reflective of divergent monocytic states in PBMCs were also evident in lung infiltrating cells (Fig. 3f). This analysis revealed 3 groups of cytokine genes that were upregulated in subsets of ETA classical monocytes. One group consisted of *CXCL8* (IL-8), *CXCL2, CCL20, CXCL3, CCL3* and *CCL4* (MIP-1β); a second consisted of *CCL2*, *CCL7*, and *IL1B* (IL-1β); and a third consisted of *IFI6*, *IFI27*, *IFI44L*, *CXCL10* (IP-10) and *CCL8*. Notably, the clusters of genes including *CCL4* and *CXCL3* were components of the Mono_cells_PDE6H gene module in peripheral monocytes, while the interferon response genes were connected to the Mono_cells_NOC2L gene module in such cells. We note that both classical and intermediate monocytes in COVID-19 patients expressed higher levels of the homing receptor CCR5 which could facilitate their trafficking from the periphery into the lung (Fig. S4a-b). Thus, the bifurcated genomic states of monocytes in peripheral blood samples from COVID-19 patients are also observed in such cells derived from ETA samples, suggesting that the divergent states reflect distinct responses to systemic infection and inflammation.

### Immune cell gene modules are predictive of COVID-19 mortality

Following our discovery that distinctive inflammatory monocyte profiles are part of immune cell genomic states that correlate with COVID-19 outcome, we next sought to determine whether a multivariate signature based on gene module scores of immune cell states on day 1 in the ICU are predictive of mortality. To evaluate whether there is a robust multivariate gene module signature associated with 90-day mortality, we first utilized the 56 significant gene module scores on day 1 to perform a PCA (Fig 4a). We found that these down-selected modules stratified the patients who died from those that survived using PC1 and PC2 (Fig. 4a). The separation between outcome evident in the first two PCs suggested that gene modules could be predictive of mortality. We next performed leave-one-out cross-validation (LOOCV) to evaluate the predictive power of this approach with data held out. In a LOOCV framework, using either a random forest or support vector machine classifier with a linear kernel on the down-selected gene modules (with down-selection done separately within each fold to avoid signal leakage) yielded accurate stratification of the patients who survived versus those that died (Methods). These machine learning approaches demonstrated area under the ROC curve scores of 0.78 and 0.88 for random forest and the support vector machine (Fig. 4b-c), respectively, suggesting that gene module scores on day 1 are predictive of eventual outcome.

**Figure 4.**
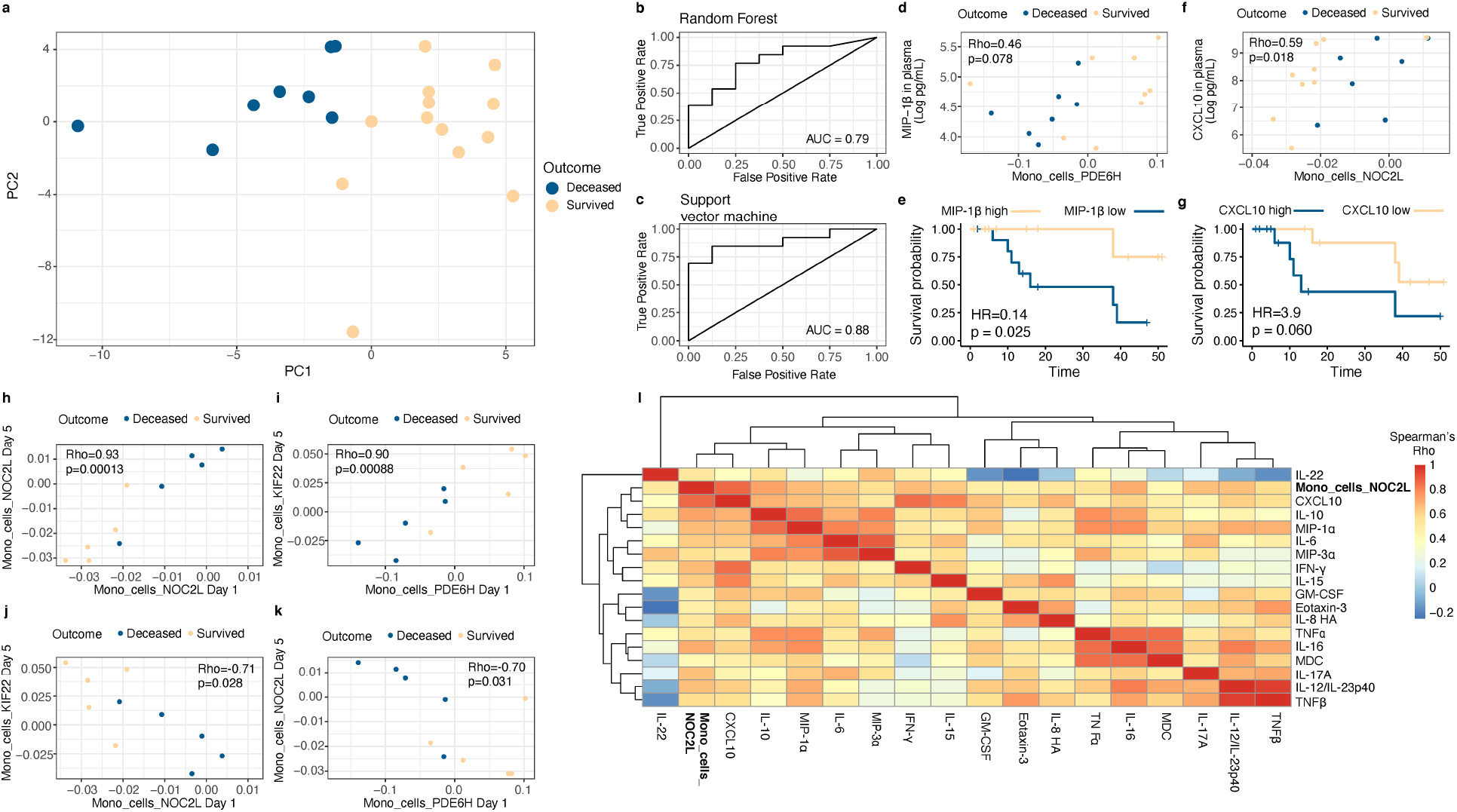
Monocyte gene modules and associated cytokines are predictive of COVID-19 mortality. a) PCA of COVID-19 patients and their outcomes based on down-selected immune gene modules scores derived from scRNAseq of PBMC samples (day 1 ICU). b-c) Area under the curve (AUC) scores derived from leave-one-out cross-validation using random forest (b) or support vector machine (c) algorithms. The diagonal line in each AUC plot represents a model with no predictive power (i.e. a 50% probability of death or survival). Random forest and support vector machine models achieved AUC scores of 0.79 and 0.88, respectively. d) Correlation plot of Mono_cells_PDE6H gene module scores with levels of MIP-1β in plasma on day 1. e) Proportional hazards regression analysis of MIP-1β levels on day 1 and mortality. f) Correlation plot of Mono_cells_NOC2L gene module scores with levels of CXCL10 in plasma on day 1. g) Proportional hazards regression analysis of CXCL10 levels on day 1 and mortality. h-k) Correlation plots of indicated monocyte modules using day 1 and day 5 scores. Mortality outcomes of COVID-19 patients analyzed by such longitudinal sampling are indicated by colored dots. I) Heatmap displaying correlation analysis of Mono_cells_NOC2L gene module scores with a network of inflammatory cytokines in plasma on day 5 in COVID-19 samples.

Given our findings that bifurcated monocyte inflammatory states may play an important role in COVID-19 patient outcome, we next evaluated whether the Mono_cells_NOC2L and Mono_cells_PDE6H gene module scores were correlated with levels of inflammatory cytokines in plasma (Methods). Briefly, we measured levels of 44 cytokines on days 1, 5 and 10 from COVID-19 patients using the Meso Scale Discovery platform (Methods). The Mono_cells_PDE6H module trended towards a significant correlation with MIP-1β in plasma (Fig. 4d), consistent with the inclusion of the gene for MIP-1β (*CCL4*) in this module. We next evaluated the extent to which this chemokine was associated with survival. Using Cox proportional hazards regression (Methods), we found that higher levels of plasma MIP-1β on day 1 were associated with improved survival (Fig. 4e). Conversely, the Mono_cells_NOC2L module correlated with CXCL10 in plasma on day 1 (Fig 4f), consistent with CXCL10 being strongly associated with response to interferon in monocytes and suggesting that the Mono_cells_NOC2L module is reflective of an interferon response in monocytes. We also found that higher levels of CXCL10 in plasma on day 1 trended towards an association with greater risk of death (Fig. 4g). The survival trends associated with these cytokines demonstrate putative biological links with the monocyte gene modules. Therefore, the monocyte gene modules capture key aspects of divergent systemic inflammatory states in severe COVID-19 disease that are predictive of mortality.

### Stable inflammatory monocyte state that portends immune sequelae and outcome

We next used longitudinal data from COVID-19 patient samples to investigate the stability of the initial monocytic cell states (day 1 ICU) and their temporal relationships with inflammatory cytokines. We first used a rank sum test to determine if monocyte gene modules were associated with outcome on days 5 and 10, and found that there were 60 monocyte modules associated with outcome on day 5 but none on day 10 (using a two-sided alpha of 10% as a cutoff; Supp. Data S3). We next evaluated which monocyte modules on day 5 were correlated with Mono_cells_NOC2L or Mono_cells_PDE6H monocyte modules on day 1 (Supp. Data S5). This analysis revealed the Mono_cells_NOC2L module to be strongly correlated with itself at days 1 and 5 (Fig. 4h) whereas the Mono_cells_PDE6H module was positively correlated with Mono_cells_KIF22 (Fig. 4i). Importantly, there was a strong negative correlation between Mono_cells_NOC2L on day 1 and Mono_cells_KIF22 on day 5 (Fig. 4j) as well as Mono_cells_NOC2L on day 5 with Mono_cells_PDE6H day 1 (Fig. 4k). These data suggest that the inflammatory monocyte state demarcated by the NOC2L module, predictive of mortality, is stable over a 5-day period in the ICU. This monocytic state can be distinguished from one delineated by the PDE6H module at day 1 that undergoes a temporal dynamic associated with survival.

Finally, we sought to evaluate the relationships of the Mono_cells_NOC2L module on day 5 was with the various inflammatory cytokines. To address this question, we correlated Mono_cells_NOC2L module scores on day 5 with levels of cytokines in plasma on day 5 (Fig. 4l). This analysis revealed that Mono_cells_NOC2L was strongly correlated with inflammatory cytokines such as CXCL10, IL-6, TNFα and IL-8 on day 5. Taken together, these data suggest that sustained viral sensing and interferon signaling by monocytes, inferred by high Mono_cells_NOC2L module scores that are strongly correlated at days 1 and 5 of ICU enrollment, may promote an inflammatory immune state involving a network of cytokines leading to immunopathogenesis and organ damage.

## Discussion

In this study, we sought to evaluate cellular and molecular features associated with the immunopathogenesis of severe COVID-19 illness that in turn were predictive of mortality (summarized in Fig. S5). This systems immunology approach involved high-dimensional longitudinal phenotyping of PBMC and cytokines of blood samples from critically ill COVID-19 patients and lower respiratory tract specimens (when possible). In agreement with other reports, we found that COVID-19 patients tended to have higher frequencies of plasmablasts^24,28^ and inflammatory monocytes^29^ and lower frequencies of CD4+ T cells^24^. Concomitant expansion of intermediate monocytes and plasmablasts have been noted in other viral diseases, such as Dengue virus^30^. In this context, the inflammatory monocytes have been shown to promote the differentiation of naïve B cells into plasmablasts which can be the source of protective or pathogenic antibodies. In this regard we note that severe COVID-19 has been associated with various types of auto-antibodies^31–33^.

Our study highlights the importance of analyzing the initial immune state of COVID-19 patients early in their ICU course by deconvolution of genomic states of peripheral immune cells based on gene modules. Earlier work in non-COVID ARDS has defined monocyte signatures in PBMC that are indicative of progression to ARDS^34^; our work demonstrates distinctive monocyte gene modules as part of a multivariate peripheral immune system state that is predictive of severe COVID-19 disease mortality. Among the gene modules on day 1 that are predictive of outcome, we uncovered a striking bifurcation of monocyte inflammatory modules. Viral sensing and interferon response genes were reflective of a monocytic state (NOC2L module) that was sustained over 5 days in the ICU and correlated with serum cytokines CXCL10, IL-6, TNFα, and IL-8. CXCL10 has been associated with severe COVID-19^15^ and IL-6, TNFα and IL-8 have been associated with increased risk of death from COVID-19^16^. In contrast, a divergent inflammatory monocytic state (PDE6H module) involving expression of the chemokine genes *CCL3* (MIP-1α)*, CCL4* (MIP-1β)*, CXCL1,* and *CXCL2* was associated with survival. Importantly, monocytic cells manifesting both of these divergent states were detected in ETA samples of severe COVID-19 patients, highlighting the relevance of analyzing PBMC.

We propose that the pathogenic viral sensing and IFN signaling monocytic state may be induced by virus entry. Intriguingly, our flow cytometry analysis identified increased expression of the SARS-Cov-2 entry receptor ACE2^35,36^ in classical and intermediate monocytes in PBMC, and increased expression of the entry cofactor NRP1^37,38^ in intermediate monocytes, suggesting that these cells may be permissive to viral entry and possibly replication, especially within the lung. We were not able to detect viral transcripts in monocytic cells of COVID-19 patients possibly due to low numbers of such transcripts resulting from non-productive infection. Even if the virus cannot replicate, after entry through ACE2 or through opsonization it could trigger innate viral sensors such as RIG-I in monocytes.

Studies of Middle East Respiratory Syndrome (MERS) Coronavirus have demonstrated infection of monocyte-derived macrophages with subsequent secretion of inflammatory cytokines such as CXCL10 and CXCL8 (IL-8)^39^. Conversely, SARS-COV abortively infects human macrophages, but triggers production of CXCL10 and CCL2^40,41^. Consistent with our findings of activated monocytes with induced expression of viral sensing and IFN response genes, an earlier study has shown that lower respiratory tract myeloid cells can harbor SARS-COV-2 and display an inflammatory phenotype^42^. Furthermore, recent work demonstrated that infected monocytes in bronchoalveolar lavage samples from patients with COVID-19 participate in a positive feedback loop in which infected myeloid cells produce T cell chemoattractants, recruiting T cells into the lung^43^. These T cells then secrete IFN-γ, contributing to release of inflammatory cytokines from alveolar macrophages, thereby promoting further T cell activation^43^. Importantly, both this emergent work and our current findings invoke activated monocytes that have sensed virus and instigate persistent alveolar inflammation. We further demonstrate that this monocyte inflammatory state is predictive of mortality in critically ill COVID-19 patients.

Type I IFN signaling has been proposed to play both a protective and pathogenic role in COVID-19 disease depending on its kinetics^20–22^. It is essential early on in infection in moderating COVID-19 disease, as patients with either IFN autoantibodies^18^ or inborn errors of type I interferon^19^ production have a much higher risk of severe COVID-19. However dysregulated interferon signaling at later times during COVID-19 disease progression appears to be pathogenic^44,45^.

The finding that *CCL4* (MIP-1β) was associated with survival is consistent with its positive prognostic role in dengue infection^46^. In hepatitis C infection, higher MIP-1β was associated with viral control following treatment with antiviral therapy^47^. Interestingly, MIP-1β is a type I interferon dependent gene but does not directly inhibit viral replication. Instead, it promotes the recruitment of monocytes to infected tissue to prevent viral spread throughout the tissue^48^. In this context, our findings suggest a model in which preferential expression of MIP-1β in relation to other IFN response genes by monocytes protects infected tissue in the lung during severe COVID-19.

High levels of inflammatory cytokines are associated with increased risk of death in COVID-19 patients^16^, but the immunologic drivers of these cytokine levels remain incompletely understood. Our analysis revealed that monocyte gene modules on day 1 dictate distinct disease trajectories, with the high levels of the interferon response associated Mono_cells_NOC2L module on day 1 correlating with its levels on day 5 in individual patients. Additionally, the Mono_cells_NOC2L module on day 5 is correlated with numerous cytokines associated with disease severity and death, including CXCL10, IL-6, TNFα and IL-8. Conversely, the high levels of the favorable monocyte state on day 1 (Mono_cells_PDE6H) was negatively correlated with Mono_cells_NOC2L on day 5, suggesting that these initially bifurcated states lead to divergent immune system dynamics. Thus, we infer that protracted interferon signaling (as reflected by consistently high Mono_cells_NOC2L module scores, ICU day 1-5) promote an immune state with correlated expression of multiple inflammatory cytokines that contributes to immunopathogenesis, organ damage, and death.

Our data regarding the temporal evolution of severe disease and the multi-faceted nature of inflammatory cytokines has important implications for COVID-19 treatment strategies. Our results suggest that the correlated expression of multiple inflammatory cytokines may partially explain conflicting results from trials of IL-6 blockade^11,49,50^, and highlight that IL-6 blockade is likely to be most beneficial early in the course of disease before multiple inflammatory cytokines become elevated. Recently, evidence from a mouse model^51^ and observational studies of rheumatic^52,53^ and inflammatory bowel disease^54^ have suggested that COVID-19 patients may benefit from anti-TNF therapy. Our results suggest that selective intervention based on new molecular diagnostics informed by our analysis of divergent inflammatory monocytic states could attenuate the coordinated elevation of multiple inflammatory cytokines and subsequent tissue damage and death. Our study adds important details of immune pathogenesis in severely ill COVID-19 patients to what is known across the spectrum of disease. Limitations of our study include the relatively small sample size of our COVID-19 patient cohort, precluding in depth analysis of relationships between flow cytometry measurements, transcriptional states of immune cells and cytokines with clinical covariates such as diabetes, age and biological sex. These covariates are important contributors to outcome and need to be more fully integrated with the high-dimensional immune system analyses in larger studies. Furthermore, emergent datasets consisting of critically ill patients that have been phenotyped as densely as our patient cohort will facilitate external validation of the findings presented here. Nevertheless, our study of severely ill COVID-19 patients admitted to the ICU has uncovered a striking bifurcation in inflammatory monocyte genomic states that is predictive of mortality outcome. These bifurcated monocytic states manifest differing temporal dynamics and are in turn linked to distinct inflammatory cytokines. Taken together, our findings may facilitate discovery of new diagnostics and therapeutics to improve outcome in severe COVID-19.

## Acknowledgements

We thank the patients and their families for their willingness to participate in this study. We also thank the frontline clinical providers (nurses, physicians, respiratory therapists and research coordinators) for their assistance in collecting biospecimens from patients in the ICU. We wish to thank everyone in the Vignali (Vignali-lab.com; @Vignali_Lab) and Bruno Labs (@BcellBruno) for all their constructive comments and advice during this project. This research was supported in part by the University of Pittsburgh Center for Research Computing through the resources provided. We thank the Hillman Cytometry Facility for assistance with flow cytometry, and the Center for Vaccine Research for access to Biosafety Level 2+ Facilities. This research was funded by the Immune Transplant and Therapy Center at UPMC (to D.A.A.V., T.C.B, and A.M.), NIH P01 AI108545 (to D.A.A.V.), NIH P01 HL103455 (to A.M.), NIH P01 HL11453 (P.R, J.S.L, B.J.M.), and a Hillman Post-doctoral Fellowship for Innovative Cancer Research (to A.R.C.).

## Authorship contributions

DAAV, TCB, AM conceived and directed the project. DAAV, TCB, and AM obtained funding for the project. GDK, BM, JSL, YZ, and AM coordinated patient enrollment, clinical phenotyping and data collection, and acquisition of biospecimens. ARC, CC, FS, GL processed samples and/or performed scRNAseq library preparation. ARC and FS performed bioinformatics analysis. AS, CW and GD performed sample processing and soluble cytokine assays. AS, TB, AR, CL, SO, and SG performed flow cytometry staining and analysis. SK, TCB and AR performed flow cytometry. TCB, LPA, SK, AS, and AR analyzed flow cytometry data. PR, AR, PVB, JD, HS, JSL, AM and GDK provided intellectual input on viral infections of the lung, computational genomics and/or machine learning. ARC, JD, HS, AM, TCB, and DAAV interpreted the results and wrote the manuscript. All authors edited and approved the manuscript.

## Materials and methods

### Clinical cohort

Following acquisition of written informed consent from patients or their legally authorized representatives, we enrolled 41 consecutive critically ill patients with acute hypoxemic respiratory failure and symptoms/signs suggestive of COVID-19 in a prospective, observational cohort study (University of Pittsburgh Institutional Review Board study number 20040036). Patients were hospitalized in ICUs at two hospitals (Presbyterian and Shadyside) within the University of Pittsburgh Medical Center system. All patients underwent at least one nasopharyngeal swab testing for SARS-CoV-2 qPCR (reference standard diagnosis at our Institution), which could be repeated at the discretion of the treating physicians when the first test was negative and significant clinical suspicion for COVID-19 remained. Based on SARS-CoV-2 qPCR results, 35 patients with at least one positive test were diagnosed with COVID-19 (COVID-19 group), whereas 6 patients had at least one negative SARS-CoV-2 qPCR test and were diagnosed with a non-COVID etiology of acute respiratory illness (non-COVID group). For comparisons against healthy controls, we also included a single blood biospecimen from 10 healthy donors.

From enrolled critically ill patients, we collected blood specimens upon enrollment (day 1), and then if the patients remained in the ICU, we collected follow-up blood samples on days 5 and 10 post-enrollment. From intubated patients, we also collected endotracheal aspirate (ETA) samples for profiling of the lower respiratory tract.

### Sample processing

Whole blood was drawn by venipuncture into EDTA tubes. Plasma was separated from whole blood by centrifugation at 400xg for 5 mins with the brake off. Following removal of plasma, blood was diluted with Hank’s Buffered Saline Solution (HBSS), and diluted blood was layered of Ficoll-Hypaque. Density gradient centrifugation was performed by spinning at 400xg for 20 minutes with the brake turned off, and the PBMC layer was removed. PBMC were then washed twice with HBSS, and carry-over red blood cells were lysed using BD Pharm Lyse per the manufacturer’s instructions. Viable cells were counted using a Nexcelom Cellometer with acridine orange and propidium iodide. Endotracheal aspirates (ETA) were collected from intubated patients in the ICU based on the condition of the patient; samples were not collected if the patient’s status was poor. ETA were processed by diluting 1.5 mL of samples up to 15 mL with HBSS, pipetting vigorously to break up aggregated sputum. Cells from ETA were then pelleted by centrifugation at 400xg for 5 minutes, and red blood cells were lysed using BD Pharm Lyse per the manufacturer’s instructions. Viable cells were counted using a Nexcelom Cellometer as described above. PBMC and ETA cells were cryopreserved in 90% FBS and 10% DMSO. Plasma was frozen at −80°C. All experiments with COVID-19 patient samples were performed in a Biosafety Level 2+ facility (with appropriate precautions) at the University of Pittsburgh’s Center for Vaccine research.

### Flow cytometry analyses

PBMC were stained for flow cytometry as previously described^25^. Briefly, cryopreserved PBMC were thawed in a water bath at 37°C, then diluted to 15 mL with warm RPMI with 10% FBS. 1-2×10^5^ cells were placed in 96 well plates and centrifuged at 400xg for 5 minutes. Supernatant was then removed, and cells were resuspened in antibody cocktails consisting of phosphate buffered saline with 10% FBS (PBS/FBS) and appropriately diluted antibodies. All antibodies were used at a 1:100 final dilution. Samples were stained for 15 minutes a 4°C, and were then washed by adding PBS/FBS and centrifuging for 5 minutes a 400xg. Viability dye in PBS (1:4000 dilution) was then added, and samples were once again incubated for 15 minutes at 4°C, followed by a subsequent wash step in PBS. Next, samples were fixed using Becton Dickinson (BD) Fix/Perm solution as per the manufacturer’s instructions. Following fixation and permabilization, intracellular antibodies were added in BD Perm/Wash solution at appropriate concentrations; samples were once again incubated at 4°C and washed. Samples were then resuspended in PBS/FBS and acquired on the appropriate flow cytometer. Samples were stained for subsequent assessment on either a Cytek Aurora 5-laser spectral flow cytometer or a Becton Dickinson 5 laser Fortessa II for standard flow cytometry. We used the following 28 antibodies::fluorophore conjugates and clones to enumerate major immune lineages using the Cytek Aurora: HLA-DR::BUV395 (BD; G46-6), CD8::BUV496 (BD; RPA-T8), CD4::BUV563 (BD; RPA-T4), CD103::BUV615 (BD; Ber-ACT8), CD45::BUV661 (BD; HI30), CD14::BUV737 (BD; M5E2), CD19::BUV805 (BD; HIB19), Ki67::BV421 (Biolegend; KI-67), FoxP3::eFluor450 (ThermoFisher; PCH101), CD38::BV480 (BD; HIT2), CD1c::PE-Cy5 (conjugated in-house; antibody from ThermoFisher, clone L161), CD45RA (Biolegend; HI100), CD62L::BV605 (Biolegend; DREG-56), CD15::BV650 (Biolegend; SSEA-1), CD25::BV711 (Biolegend, BC96), CD20::BV750 (BD; 2H7), CD141::BV785 (BD; M80), CD36::FITC (Biolegend, 5-271), CD3::Spark Blue 550 (Biolegend; SK7), CD11b::PerCP-Cy5.5 (Biolegend; LM2), CD56::PerCP-EF710 (Invitrogen, CMSSB), ACE2::PE (R&D Systems; 171606), CD16::PE-TexasRed (Biolegend; 3G8), CD27::BV510 (Biolegend; O323), CD138::APC (Biolegend; MI15), CD11c::Alexa700 (Biolegend; 3.9), and CCR2::APC-Cy7 (Biolegend::K036C2). We also used Zombie NIR fixable viability dye (Biolegend) at a 1:1000 dilution for this panel. For our monocyte specific Fortessa II panel, we used the following 14 antibodies::fluorophore conjugates and clones: HLA-DR::BUV395 (BD; G46-6), CD14::BUV737 (BD; M5E2), CD16::BV412 (Biolegend; 3G8), CCR5::BV510 (Biolegend; J418F1), CD3::BV650 (Biolegend; UCHT1), CD19::BV650 (Biolegend; UCHT), CD20::BV650 (Biolegend; 2H7), CD56::BV650 (Biolegend; 5.1H11), CD163::BV711 (Biolegend; GHI/61), CCR2::BV785 (Biolegend; K036C2), ACE2::APC (R&D Systems; 171606), CD11c::Alexa700 (Biolegend; 3.9), NRP1::PE (Biolegend 12C2), and CD36::PE-Cy7 (Biolegend; 5-271). We also used eFluor 780 Fixable Viability Dye (ThermoFisher) for this panel at a 1:4000 dilution. Flow cytometry was performed in the Hillman Cancer Center Flow Cytometry Facility.

### Soluble cytokine/chemokine quantification by Meso Scale Discovery

We utilized the V-PLEX Human Cytokine 44-plex Kit from Meso Scale Discovery (MSD) to quantify levels of chemokines and cytokines in plasma of patients with COVID-19. The following soluble markers were included: Eotaxin, Eotaxin-3, GM-CSF, IFN-γ, IL-1α, IL-1β, IL-1RA, IL-2, IL-3, IL-4, IL-5, IL-6, IL-7, IL-8, IL-8 (HA), IL-9, IL-10, IL-12/IL-23p40, IL-13, IL-15, IL-16, IL-17A, IL-17A/F, IL-17B, IL-17C, IL-17D, IL-21, IL-22, IL-23, IL-27, IL-31, IP-10, MCP-1, MCP-4, MDC, MIP-1α, MIP-1β, MIP-3α, TARC, TNFα, TNFβ, TSLP, and VEGF-A. Assays were performed as per the manufacturer’s instructions. Briefly, plasma samples were thawed and on average 25 μL of plasma samples were diluted as recommended in a MSD 96-well assay plate. Calibrators were added to wells on each plate in parallel. Plates were then sealed and incubated overnight at 4°C. Next, the plates were washed 3 times with 150 μL per well of MSD wash buffer. Following washing, 25 μL of detection antibodies was added to each well, and plates were then sealed and incubated at room temperature for 2 hours with shaking. After 2 hours, the plates were once again washed 3 times with MSD wash buffer and 150 μL of MSD Read Buffer T was added to each well and the plates were analyzed on the MESO QuickPlex SQ 120MM. Sample concentrations for each marker were then calculated based on the respective standard curve. For analysis, the limit of detection was set at 50% of the lowest limit of detection across all analytes (i.e. 0.045 pg/mL).

### Single-cell RNAseq library generation and sequencing

Single-cell RNAseq (scRNAseq) was performed using 5’ v1 kit from 10X Genomics as per the manufacturer’s instructions. Libraries were created from either fresh PBMC or were prepared from batches of cryopreserved PBMC. For fresh processing, libraries were created immediately following isolation of cells. For cryopreserved samples, samples were thawed as described above for flow cytometry. Sample multiplexing was performed using CITEseq. To achieve this, cells were first stained with TotalSeq-C antibodies for 15 minutes at 4°C, followed by washing in PBS with 10% FBS and centrifugation for 5 minutes at 400xg. Two samples (each stained with unique TotalSeq-C antibodies) were then pooled and loaded per lane of the 10X chip to permit sample multiplexing. Libraries were prepared following the manufacturer’s instructions, with the additional preparation of cell hashing libraries using the manufacturer’s protocol for Feature Barcode libraries. Final libraries concentration and size distributions were quantified using a BioAnalyzer as per the manufacturer’s instructions. Samples were sequenced on a NovaSeq at the UPMC Genome Core using a read 1 length of 28 cycles, read 2 length of 91 cycles and an i7 read length of 8 cycles.

### Generation of gene expression and feature barcode matrices

Following sequencing, NovaSeq runs were downloaded from the UPMC Genome Core to the University of Pittsburgh Center for Research Computing High Throughput Cluster. Samples were then demultiplex using bcl2fastq (Illumina), using a base mask of Y28, I8, Y91 and setting the stringency to allow no barcode mismatches. Following demultiplexing, gene expression reads were then aligned to the reference genome using CellRanger v3.1.0, and feature barcode matrices were created for each sample. Importantly, we added the reference sequence for SARS-CoV-2 (NC_045512.2) to the genome and the GTF to facilitate detection of SARS-CoV-2 transcripts. Cell hashing libraries were aligned to TotalSeq-C barcodes using CITE-seq-Count v1.4.3^55^, using a sample-specific cell barcode whitelist (i.e., only cell barcodes from cells identified by CellRanger from each sample were included in the whitelist).

### Identification of individual samples from cell hashing

After generation of gene expression and feature barcode matrices, downstream analysis was performed using Seurat v3.1.4^56^ in R 3.6.0. Gene barcode matrices and associated feature barcode matrices containing cell hash expression values were read into R. To identify individual samples, feature barcode matrices were first log-normalized using a centered log ratio transformation (CLR) implemented in Seurat. Individual samples were identified by unique expression of the anticipated TotalSeq-C antibody and the absence of alternative TotalSeq-C antibodies. To permit automated identification of samples, k-means clustering was performed on CLR normalized expression values for each sample to identify cut-offs for negative and positive TotalSeq-C counts. Samples were visualized on bi-variate x/y plots to confirm adequate cutoffs. Cells with expression levels above the cut-offs for both TotalSeq-C antibodies were considered doublets, and were excluded from downstream analysis.

### Visualization of scRNAseq data and identification of cell types

We utilized the data integration workflow^56^ provided in Seurat to integrate between fresh and cryopreserved samples. Briefly, we independently identified highly variable features between fresh and cryopreserved samples and selected 2,000 integration features from this combined set of highly variable features. Selected integration features were then independently scaled across all cells in both fresh and cryopreserved samples, and PCA was performed using the scaled expression levels in each dataset. Next, integration anchors were identified using a reciprocal PCA across the first 30 PCs. Integrated data was then used for downstream PCA, visualization and clustering. Dimensionality reduction for visualization was next performed using Uniform Manifold Approximation Embedding (UMAP)^26^ implemented in Seurat. Clusters were identified using graph-based clustering in Seurat, and inspection of canonical lineage markers and their association with each cluster was used to identify cell types. One cluster of cells expressed high levels of genes involved in the cell cycle, but included multiple cell types. Cell types were resolved from this cluster by bioinformatically isolating these cells and inspecting lineage specific markers and then applying those labels to the cells in the overall UMAP.

### Generation of gene module scores for immune lineages

To quantify gene co-express modules within each immune lineage, we first bioinformatically isolated each major immune lineage (i.e. CD4+ T cells, CD8+ T cells, B cells, plasmablasts, monocytes and NK cells) and verified there were no contaminating immune cells from other lineages present (e.g. there were no clusters expressing *MS4A1* in the CD4+ T cell data, no clusters expressing *CD14* in the CD8+ T cells, et cetera). Next, we utilized both the SCENIC^57^ R package (version 1.1.2-2) and the Arboreto package (version 0.1.0) implemented in PySCENIC^27^ in Python (Bioconda 3.7-2019.03). In R, genes were filtered based on a minimum count of 3 in 1% of cells, and expression in at least 1% of cells. Filtered expression matrices and a list of expressed transcription factors were then exported to be used in GRNBoost2 from the Aboreto package in Python. Gene modules identified by GRNBoost2 were filtered based on having greater than 20 genes and fewer than 200 genes, then scored across all cells in an expression dataset using the AddModuleScore function in Seurat. Median module scores were derived for each patient across all immune lineages, providing great than 20 cells of a particular lineage was present.

### Principal component analysis and machine learning analysis

Principal component analysis was performed using the R package irlba. Centered and scaled values of either immune cell frequencies or median gene module scores per patient derived from Arboreto were used as input. If a patient did not have a score for a module (i.e. due to insufficient cells present in that sample), the median module score across all patients for that module was used as the interpolated value. Sample embeddings and variable loadings were extracted from resulting PCA and used for visualization.

To assess the ability of gene modules to predict outcome, we utilized the R package caret^58^ to perform machine learning with cross-validation using a random forest and a support vector machine (SVM) with a linear kernel. Cross-validation was performed using a leave-one-out analysis. To prevent data leakage between folds of cross validation, gene modules from day 1 samples that had a p<0.05 for a Wilcoxon rank sum test between patients who survived versus those that died were selected within each fold. Then, modules were scaled and PCA was performed for dimensionality reduction in each fold, and PC1 and PC2 were used as predictive variables. Tuning parameters for each algorithm were selected internally within each round of cross-validation. Receiving operating curves and area under the curve were calculated using the R package pROC^59^.

### In-depth analysis of myeloid lineages

Further dissection of myeloid lineages was performed by bioinformatically isolating and re-clustering all myeloid cells. Datasets were once again integrated for downstream visualization and clustering based on fresh versus frozen status using the workflow described above. Differentially expressed genes were identified using a Wilcoxon rank sum test as implemented in Seurat. Gene set enrichment analysis was performed as previously described using the R package singleseqgset^25^. Statistically significant gene sets were identified as those that had p values corrected for false discovery rate of less than 5%.

### Statistical analyses

Survival analysis was performed using Cox’s proportional hazard regression analysis as implement in the R package survival^60,61^. Likelihood ratio tests were used to determine the statistical significance of the survival models. Wilcoxon rank sum tests were used to evaluate differences in immune cell frequencies and mean fluorescence intensity by flow cytometry, to calculate differentially expressed genes from the scRNAseq analysis, and identify gene modules that were statistically different between patients who survived and those that died. Spearman’s correlation was used to calculate the correlation between gene module scores and cytokine levels in plasma, between gene modules on day 1 and day 5, and between cytokines and gene modules on day 5. A two-sided alpha of 5% was considered significant unless otherwise noted.

## Data availability

Both raw and processed transcriptomic data will be available through the Gene Expression Omnibus database.

## Code availability

The Seurat package was used for scRNAseq normalization, scaling, dimensionality reduction, UMAP visualization, clustering, and differential gene expression analysis. Code for these steps is available through Seurat’s website (https://satijalab.org/seurat/).

Code for gene set enrichment analysis is available at www.github.com/arc85/singleseqgset. Additional code is available upon request.

## Supplementary information

Table S1. Clinical characteristics of study participants across cohorts.

Table S2. Associations between clinical covariates and outcome.

Table S3. Patient samples utilized for flow cytometry, single-cell RNAseq, and measurement of cytokine levels.

Figure S1. Gating strategy to quantify immune cell frequencies in PBMC.

Figure S2. Evaluation of immune cell frequencies by outcome in non-COVID ARDS and COVID-19 patients.

Figure S3. Description of the Arboreto pipeline for identification of transcription factor associated co-expression modules.

Figure S4. Monocytes from COVID-19 patients have higher levels of CCR5 versus healthy donors.

Figure S5. Factors associated with death versus survival in critically ill COVID-19 patients.

## Extended Data

Supplementary Data S1. Gene modules and top 200 genes from each module across all cell types.

Supplementary Data S2. Gene modules associated with differences between clinical groups.

Supplementary Data S3. Gene modules associated with differences in COVID-19 outcome.

Supplementary Data S4. Gene set enrichment analysis of classical, intermediate, and non-classical monocytes.

Supplementary Data S5. Correlations between monocyte modules on day 1 and day 5.

