## Extended Data Figures for "Bifurcated monocyte states are predictive of mortality in severe COVID-19"

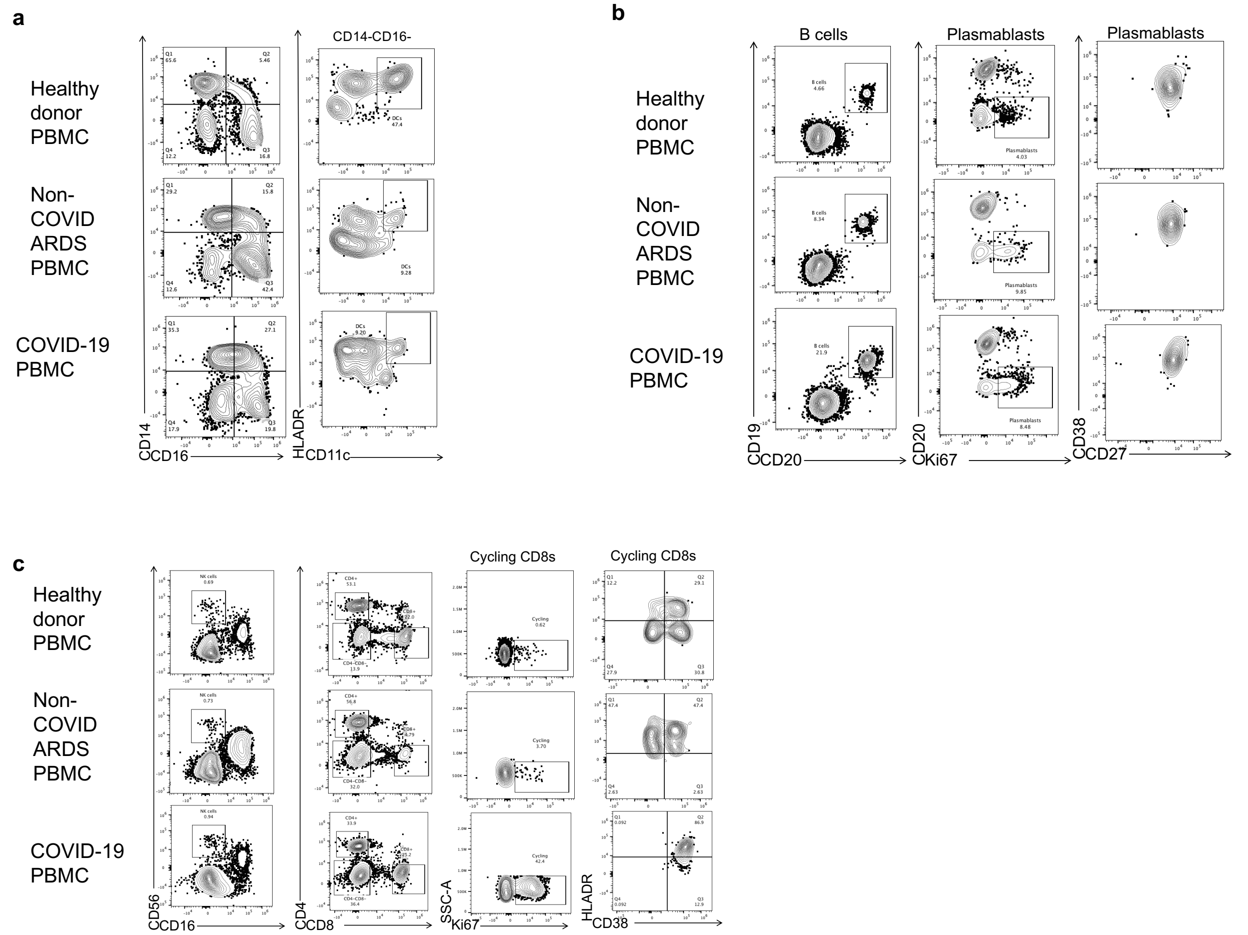


**Figure S1. Gating strategy to quantify immune cell frequencies in PBMC.** We utilized the Cytek Aurora to perform high-dimensional flow cytometry to enumerate cell frequencies in peripheral blood. All samples were first gated on forward and side scatter to identify lymphocytes based on size and granularity, followed by gating live singlet events, and then all CD45+ cells. Cell frequencies are a percent of CD45+ cells unless otherwise noted. a) Classical monocytes were gated as CD14+CD16-, intermediate monocytes were CD14+CD16+, and non-classical monocytes were CD14-CD16+. Dendritic cells were gated as CD14-CD16-CD11c+HLA-DR+ cells. b) B cells were identified as CD19+CD20+ cells, and plasmablasts were identified as CD20dimKi67+ cells. Plasmablasts also co-expressed CD38 and CD27. c) NK cells were gated as either CD56+CD16+ cells or CD56+CD16dim cells. CD4+ and CD8+ T cells were characterized by their reciprocal expression of each marker. CD8+ T cells were further sub-divided into cycling Ki67+CD8+ T cells and Ki67+CD38+HLA-DR+ cycling CD8+ T cells, and were quantified as a frequency of total CD8+ T cells.

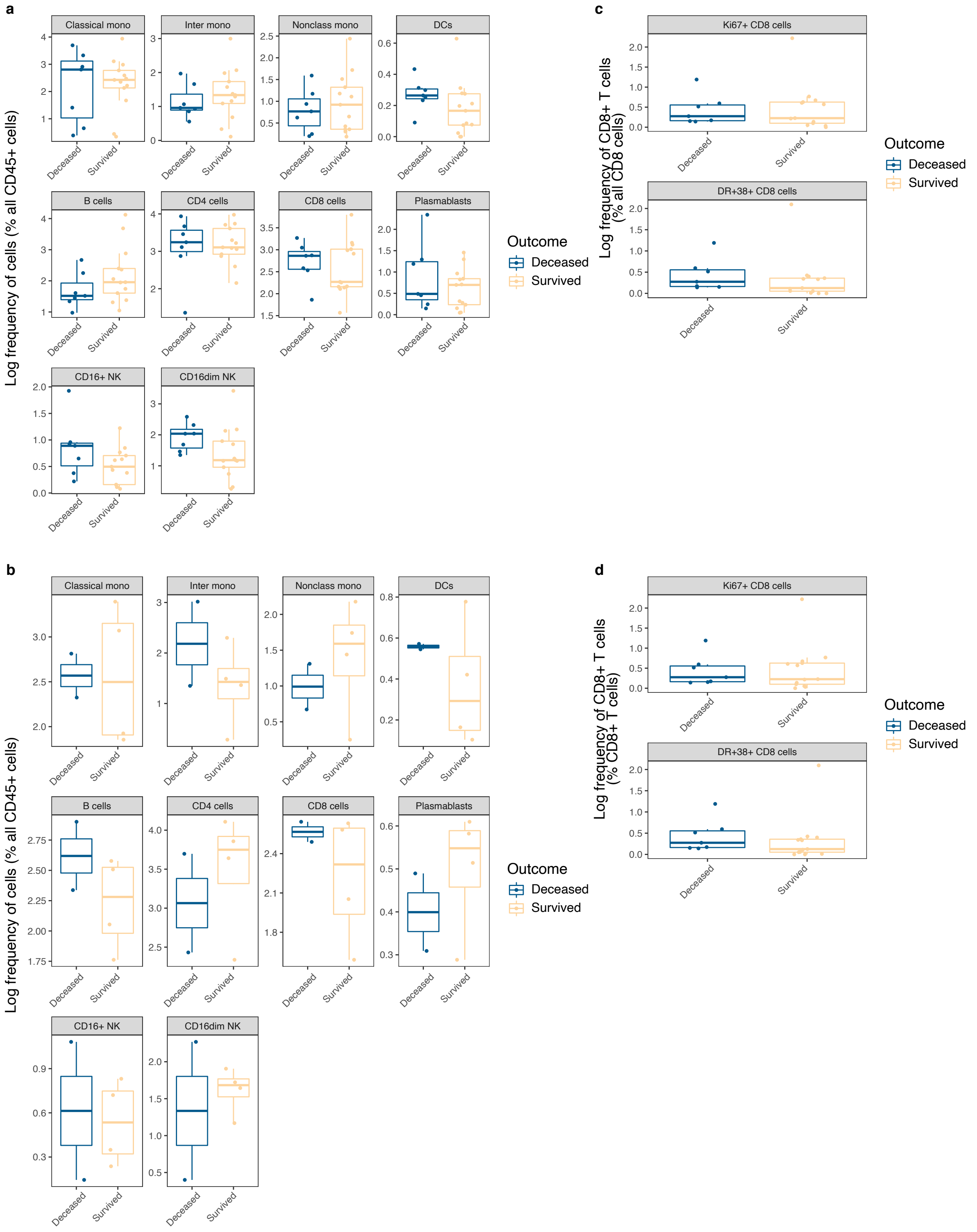


**Figure S2. Immune cell frequencies by outcome in non-COVID ARDS and COVID-19 patients.** Comparison of frequencies of immune cells on day 1 by outcome as measured by 90-day mortality. a) There were no statistically significant differences in immune cell frequencies between COVID-19 patients who died and those who survived (p values for T tests ranged from p=0.09 to p=0.86). b) There were no statistically significant differences in immune cell frequencies between non-COVID ARDS patients who died versus survived (p values ranging from p=0.30 to p=0.99). c) Comparison of all cycling CD8+ T cells and HLA-DR+CD38+ cycling CD8+ T cells between surviving and discharged COVID-19 patients revealed no significant differences. d) No statistically significant differences in mortality were evident between cycling and HLA-DR+CD38+ cycling CD8+ T cells in non-COVID ARDS patients.

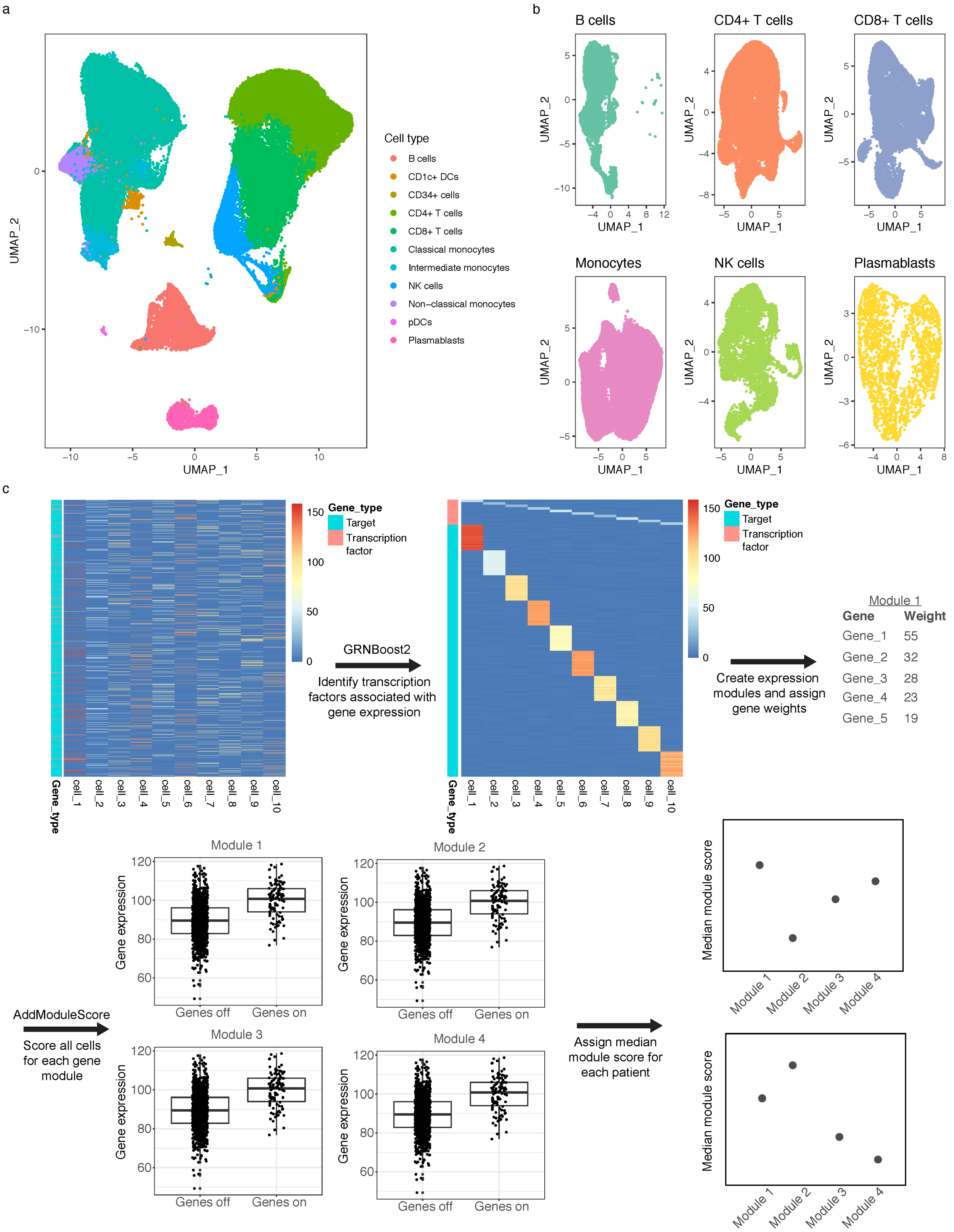


**Figure S3. Arboreto pipeline for identification of gene modules from scRNAseq.** Overview of gene module discovery using Arboreto. a) Cell types were first classified based on canonical gene expression profiles. b) Individual cell subtypes were bioinformatically isolated for downstream gene module discovery. c) Arboreto utilizes the GRNBoost2 framework to identify genes that fall under the control of given transcription factors. An expression matrix (top left) is organized into sets of genes that are co-expressed with a given transcription factor (top center) using tree-based methods, and a weight for the co-expression is assigned based on the significance from the tree-based regression. Next, each cell is assigned a score per module based on the genes within each module using AddModuleScore from Seurat. Finally, median module scores per patient can be used to capture the overall influence of a gene module in a given patient.

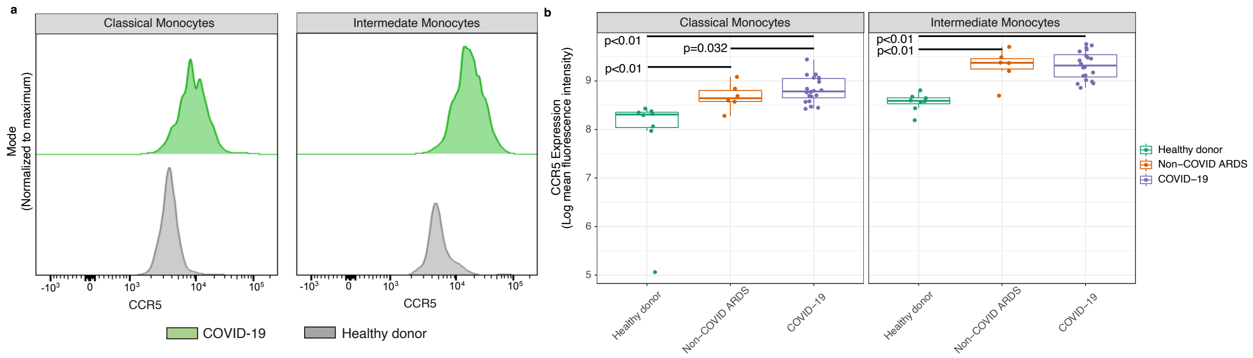


**Figure S4. Higher levels of CCR5 in non-COVID ARDS and COVID-19 patients versus healthy donors.** a) Representative flow cytometry plots of CCR5 expression. b) Quantification of CCR5 mean fluorescence intensity across patient cohorts for classical and intermediate monocytes.

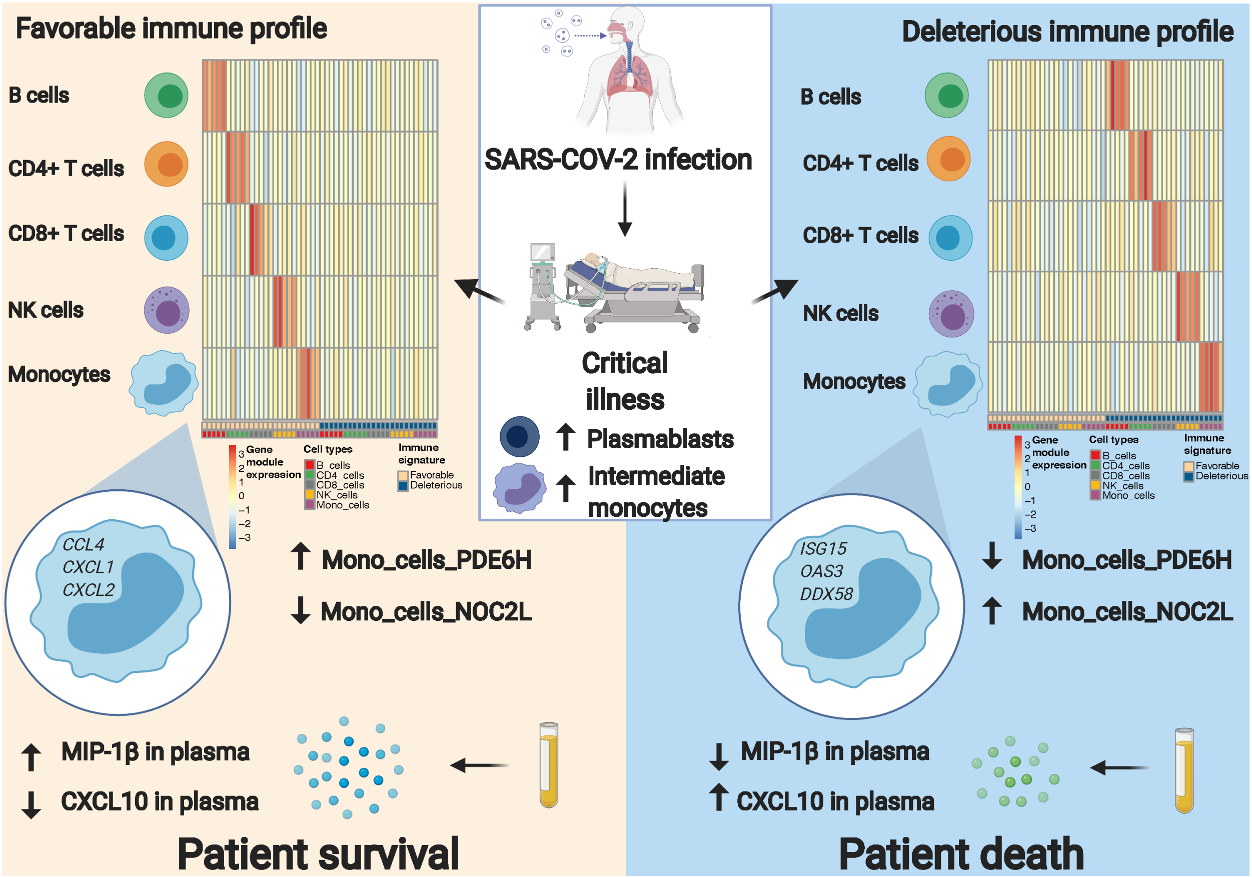


**Figure S5. Factors associated with death versus survival in critically ill COVID-19 patients.** Patients with COVID-19 have distinct transcriptional immune profiles on day 1 in peripheral blood that are predictive of eventual death or survival. In particular, a bifurcated monocyte state characterized by several chemokines including *CCL4* was associated with survival whereas an interferon signaling and virus sensing profile was associated with death. Further, higher plasma levels of MIP-1β (CCL4) were associated with improved survival, while higher plasma levels of CXCL10 were associated with death. Parts of this figure were created using BioRender.com.
